# Convergent reductive evolution in bee-associated lactic acid bacteria

**DOI:** 10.1101/2024.06.28.601270

**Authors:** Ana Pontes, Marie-Claire Harrison, Antonis Rokas, Carla Gonçalves

## Abstract

Distantly related organisms may evolve similar traits when exposed to similar environments or engaging in certain lifestyles.

Several members of the Lactobacillaceae (LAB) family are frequently isolated from the floral niche, mostly from bees and flowers. In some floral LAB species (henceforth referred to as bee-associated), distinctive genomic (e.g., genome reduction) and phenotypic (e.g., preference for fructose over glucose or fructophily) features were recently documented. These features are found across distantly related species, raising the hypothesis that specific genomic and phenotypic traits evolved convergently during adaptation to the floral environment.

To test this hypothesis, we examined representative genomes of 369 species of bee-associated and non-bee-associated LAB. Phylogenomic analysis unveiled seven independent ecological shifts towards the floral niche in LAB. In these bee-associated LAB, we observed pervasive, significant reductions of genome size, gene repertoire, and GC content.

Using machine leaning, we could distinguish bee-associated from non-bee-associated species with 94% accuracy, based on the absence of genes involved in metabolism, osmotic stress, or DNA repair. Moreover, we found that the most important genes for the machine learning classifier were seemingly lost, independently, in multiple bee-associated lineages. One of these genes, *adhE*, encodes a bifunctional aldehyde-alcohol dehydrogenase associated with the evolution of fructophily, a rare phenotypic trait that was recently identified in many floral LAB species. These results suggest that the independent evolution of distinctive phenotypes in bee-associated LAB has been largely driven by independent loss of the same set of genes.

**Importance:** Several lactic acid bacteria (LAB) species are intimately associated with bees and exhibit unique biochemical properties with potential for food applications and honeybee health. Using a machine-learning based approach, our study shows that adaptation of LAB to the bee environment was accompanied by a distinctive genomic trajectory deeply shaped by gene loss. Several of these gene losses occurred independently in distantly related species and are linked to some of their unique biotechnologically relevant traits, such as the preference of fructose over glucose (fructophily). This study underscores the potential of machine learning in identifying fingerprints of adaptation and detecting instances of convergent evolution. Furthermore, it sheds light onto the genomic and phenotypic particularities of bee-associated bacteria, thereby deepening the understanding of their positive impact on honeybee health.

## Introduction

Ecological lifestyles shape the content of microbial genomes. For instance, microorganisms that engage in strictly dependent relationships with their hosts, such as parasites or intracellular symbionts, generally undergo major reduction in genome size and gene repertoire (1, 2). Numerous examples have been found across microorganisms such as fungi and bacteria. For instance, species belonging to the fungal lineage Microsporidia, strictly composed of obligate intracellular parasites, are devoid of many central pathways involving carbohydrate and amino acid metabolism, exhibiting some of the most compact eukaryotic genomes (3–5). In bacteria, genome reduction associated symbiotic lifestyles has been identified in independent lineages from diverse bacterial groups (6, 7). In addition to reduced gene repertoires, symbiotic bacterial genomes sometimes exhibit other distinctive features such as rapid sequence evolution, codon reassignments or low GC content (6). While most losses are possibly the result of relaxed selection acting on functions that can be supplied by the host, such as those involved in metabolic processes, others might be the result of genetic drift during the shift to a more confined and restricted habitat, where the effect of small effective population result in reduced opportunities for gene exchange and homologous recombination, resulting in strong intergenerational bottlenecks (1, 6, 8–10).

Other gene losses can however be adaptive when organisms are adjusting to a new environment (11). For instance, loss of function mutations in genes involved in multiple pathways (e.g., DNA repair or adhesion to neutrophils) may increase pathogenicity or drug resistance in bacterial species, providing an selective advantage for survival in the human body (12).

Recently, a unique group of lactic acid bacteria (LAB) belonging to *Fructobacillus* and *Lactobacillus* genera was described based on their unusual metabolic characteristics and their distinctive ecological association with fructose-rich environments involving flowers, fruits, and flower-visiting insects, such as bees (13–15). Many of these bacteria have been described as bee symbionts (16, 17), however they are found across the floral niche, suggesting that they inhabit the bee gut but can thrive outside their hosts. Adaptation to this fructose-rich environment is thought to have promoted a major metabolic remodelling towards efficient fructose utilization, because these species use fructose more efficiently than glucose (15, 18). This unusual metabolic feature, dubbed as fructophily, is seemingly associated with the partial or complete loss of a gene encoding a bifunctional acetaldehyde-alcohol dehydrogenase (*adhE*), rendering these species unable to engage in alcoholic fermentation (18–20). Many fructophilic LAB (FLAB) underwent major losses of genes involved, for instance, in carbohydrate metabolism (such as phosphotransferase system, PTS) (16). Whether these losses result from adaptation to the floral environment, from the symbiotic relationship with bees or both remains elusive.

Hence, we set out to look for evidence of convergent evolution in floral bacteria (henceforth referred to as bee-associated) across the entire Lactobacillaceae family (369 species; henceforth referred to as LAB), which includes four distinct clades of bee-associated species (21, 22). We used a phylogenomic framework onto which ecology and genomic information was mapped. We inferred that an association with bees evolved multiple (seven) times independently in LAB, involving entire clades (four) and single species (three). Supporting previous reports, we observed that genome reduction is pervasive across bee-associated LAB and found other distinctive features of symbiont genomes such as low GC content. Moreover, we observed that absence of *adhE* is a distinctive feature of bee-associated LAB. Although phenotypic information regarding fructophilic behaviour is missing for a significant portion of the species under study, the absence of *adhE* in bee-associated LAB indicates that loss of alcoholic fermentation and evolution of fructophily might be strongly linked to adaptation to this environment. To find additional genomic fingerprints of convergent evolution, we used a machine learning algorithm to show that association with the floral environment can be predicted from functional genomic data with very high (94%) accuracy. The *adhE* gene was the most important predictor; however, prediction accuracy remained unaltered when we removed *adhE* from the dataset, indicating that other genomic features are contributing. These features included genes involved in carbohydrate and amino acid metabolism, osmotic stress, and DNA repair, which are distinctively absent from bee-associated LAB. Moreover, we found that some of these genes were likely lost in the branches leading to distantly related bee-associated clades, suggesting that loss of the same genes occurred independently in association with adaptation to the bee environment. These results revealed that convergent adaptation to the bee environment in LAB was seemingly accomplished through highly similar evolutionary mechanisms, some of which (i.e., loss of AdhE) are associated with distinctive metabolic features (i.e., fructophily).

## Results

### Multiple instances of ecological association with the bee environment across Lactic Acid bacteria (LAB)

To ascertain the distribution of bee association across lactic acid bacteria, we retrieved all representative genomes from species belonging to the Lactobacillaceae (LAB) family (369, as of 19^th^ January 2023) (Table S1). We specifically focused on LAB because multiple species were previously documented to be frequently isolated from the floral environment (13, 14, 20). We inferred a phylogenomic tree using 180 single copy orthogroups present in at least 50% of species as obtained from OrthoFinder (23). The resulting species tree (Figure 1, Figure S1) recapitulates the phylogenetic relationships among the main lineages within LAB (21).

**Figure 1.**
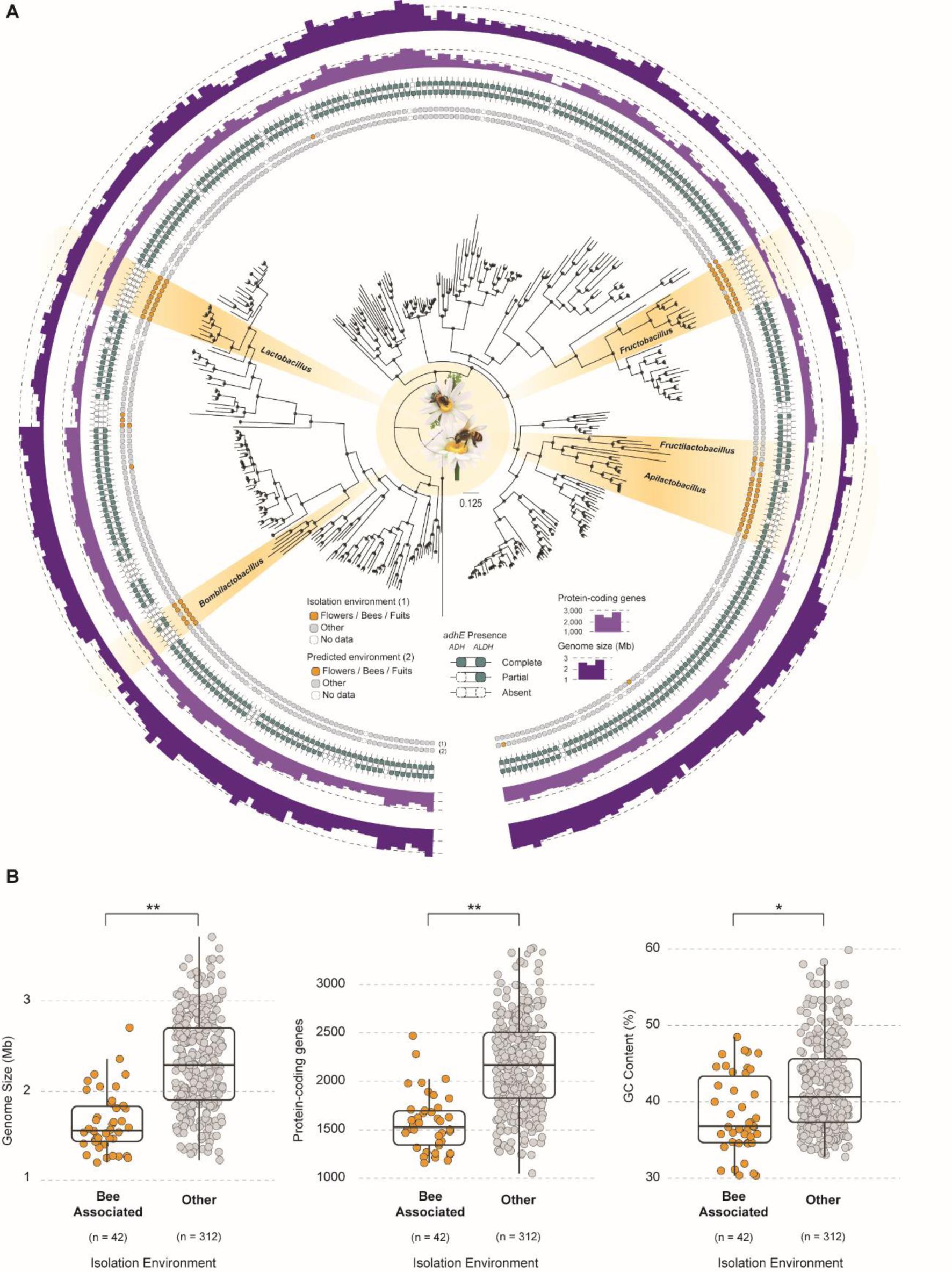
**Phylogenomic tree of the Lactobacillaceae**. (A) Maximum likelihood phylogenomic tree comprising 369 LAB species inferred from the concatenated alignment of 180 single copy orthogroups (SCO) and rooted with *Lactococcus lactis* and *Enterococcus massiliensis*. The isolation source and predicted environment are depicted in the first two rings, respectively. Presence/absence of *adhE*, number of protein-coding genes, and genome size are also shown in the other three rings. (B) Boxplots depicting comparisons between bee-associated species and others regarding genome size, number of protein-coding genes, and GC content.

We next inspected ecological association with the floral environment (bees, flower, and fruits; henceforth referred to as the bee environment) across the entire dataset. Ecological association was established according to the substrate of isolation of the strain for which the genome sequence was obtained. Most of the 369 species were isolated from anthropic environments (processed foods) and the gut of mammals (Table S1), however, 42 species were isolated from bee-associated environments. We identified four distinct instances of bee association involving entire clades (Figure 1, Figure S1): *Bombilactobacillus, Fructobacillus*, *Fructilactobacillus* - *Apilactobacillus* genera and a subclade within the *Lactobacillus* genus; and three instances of floral association in distantly related single species: *Companilactobacillus musae*, *Holzapfelia floricola*, and *Secundilactobacillus yichangensis*. Most species (∼60%) were isolated from bees while the remaining 40% were isolated from flowers, fruits, or honey (Table S1).

Using this classification, we can infer that seven independent ecological shifts towards bee-associated environments likely occurred within LAB. The clades identified include species described in previous studies as being fructophilic lactic acid bacteria (FLAB) (15, 21).

### Genome reduction is pervasive across bee-associated LAB

Distinctive genome reduction was previously reported for several FLAB isolated from the floral niche (21). We therefore assessed the genome sizes of 369 species belonging to LAB. We observed that genome size considerably varies across the family (1.2 – 3.7 Mb). When comparing bee-associated species with all the others, and accounting for phylogenetic relatedness (24), we observed that bee-associated genomes are significantly smaller (*p-value* = 0.001, phylANOVA). In line with genome reduction, we also observed that bee-associated species encode a significantly lower number of protein-coding genes (*p-value* = 0.001, phylANOVA), indicating that genome reduction is associated with gene loss (Figure 1A, 1B). In addition to decreased genome size, symbiotic bacteria can also exhibit significantly lower GC content when compared to their non-symbiotic relatives (6). By analysing GC content across LAB, we observed that bee-associated species present significantly lower GC content than non-bee-associated species (*p-value* = 0.046, phylANOVA) (Figure 1B).

One of the most distinctive known features of bee-associated FLAB is the absence of the *adhE* gene, encoding a bifunctional acetaldehyde/alcohol dehydrogenase, which is linked to the evolution of fructophily in several *Fructobacillus* and *Apilactobacillus* species (16, 21, 25). To ascertain whether the loss (total or partial) of this gene is more broadly associated with species thriving in the bee environment, we looked for the presence of AdhE across the entire dataset using an HMM-based sequence similarity approach (26). We found that 80% of bee-associated species either partially (ethanol dehydrogenase domain absent) or completely lack AdhE (Figure 1, Figure S1). A phylogenetically informed comparison of the distributions of niche (bee and other) and AdhE (present and absent) showed that the pattern of occurrence of AdhE is statistically dependent of the niche (*p-value* = 0.0008, Chi-squared test).

Although phenotypic information regarding fructophily is limited, by inspecting the available literature we found that at least 50% of the bee-associated LAB that lack Adh activity have been described as fructophilic (Figure S1) (14, 15, 18, 20, 21, 27). This suggests a link between loss of alcohol dehydrogenase function and adaptation to the bee environment in LAB. Interestingly, even in clades where loss of AdhE is pervasive, some species encode apparently functional AdhE enzymes. For instance, in the *Apilactobacillus*-*Fructilactobacillus* clade, some species totally lack AdhE (2), others only lack the Adh domain (10), and some have apparently functional enzymes (7). Since these three patterns do not strictly align with the phylogenetic relationships between these species, it seems plausible that loss of Adh activity happened multiple times in this niche, both in the most recent common ancestor (MRCA) of entire clades but also in single species and can happen in a stepwise manner.

### Machine learning analyses reveal that absence of specific genes accurately predicts bee association in LAB

Although total or partial absence of AdhE seems to be common across bee-associated LAB, we next inspected whether other genes are also fingerprints of adaptation to this environment. For that, we performed genome wide functional annotations (KEGG) for all species under study and employed a machine learning approach. Species for which a low number of proteins were mapped to a functional KEGG annotation were excluded (ratio of protein-coding genes / KEGG annotated proteins < 3, Figure S2). In this way, KEGG annotations (presence/absence) and ecological information for 360 species were submitted to a trained and supervised random forest (RF) classifier (28–30). Species were classified according to their ecology in the following manner: 1–isolated from bees, flowers, or fruits; 0–not isolated from bees, flowers, or fruits (other); NA–no information (Table S1).

We obtained highly accurate predictions (balanced accuracy of 94%) of niche association (Figure 2A). To determine which features are contributing to the RF classification, we investigated the 20 most important features (Figure 2B). In line with the evidence of genome reduction, we found that the most important predictive genes are less prevalent in bee-associated species when compared to non-bee-associated species (Figure 2C, p-value < 2.2e^-16^, Chi- squared test). The top predictive feature was the *adhE* gene (K04072), followed by *glxK/garK* (K00865), and *ohyA* (K10254). The loss of *adhE* has been previously associated with FLAB, however absence of *glxK/garK*, encoding a glycerate 2-kinase, and *ohyA*, encoding an oleate dehydratase, have not been previously reported to our knowledge. *glxK/garK* is involved in the glyoxylate and glucarate/galactarate utilization pathways, and *ohyA* is possibly associated with the conversion of unsaturated fatty acids and might play a role in the evasion of the human host innate immune response by *Staphylococcus aureus,* through inactivation of antimicrobial unsaturated fatty acids (31).

**Figure 2.**
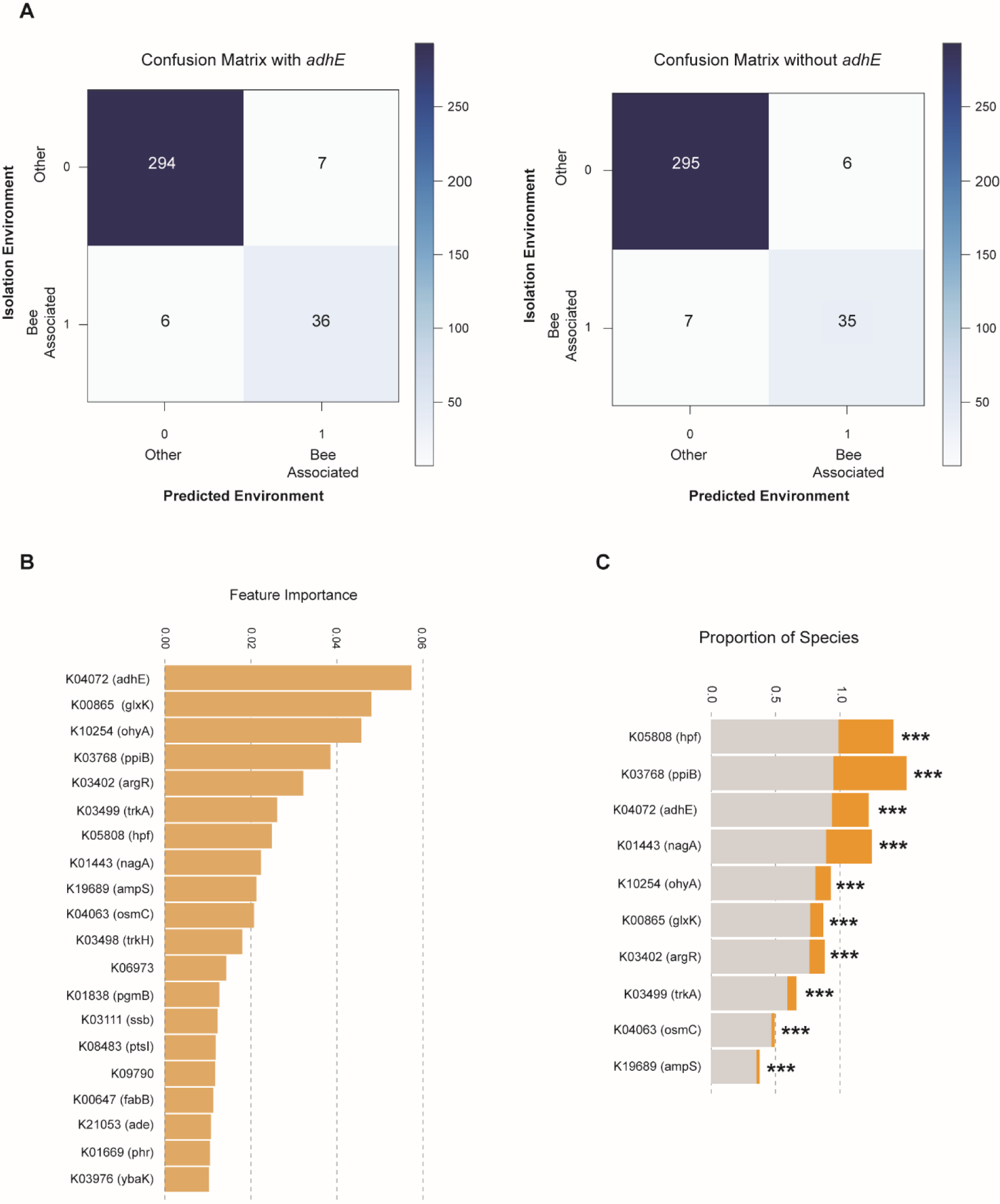
Bee-associated LAB can be predicted from genomic data with high accuracy. (A) Confusion matrix with and without AdhE (left and right, respectively) across 343 species of the Lactobacillaceae family. (B) Top 20 most important features for the RF classifier. (C) Distribution of the top 10 most important KEGG features across species belonging to bee-associated (orange) and non-bee-associated (grey) groups. Bar plots represent the proportion of species in which the KEGG annotation was found to be present. Statistically significant differences between the proportion of species (bee-associated and others) encoding each function are annotated next to the respective KEGG (Chi-squared test).

Since classification accuracy is being driven by absence of genes, we next confirmed that functional annotation issues did not influence our results. For that, we searched for the *ohyA* gene using an HMM-based approach across LAB proteomes and confirmed its absence in bee-associated *Fructobacillus*, *Apillactobacillus*, and in the bee-associated *Lactobacillus* subclade (Figure S1, Table S1). Because of its importance for the RF classifier and of its notable absence in bee-associated species, we next determined its distribution across LAB species. Mapping the presence/absence of *ohyA* to the species phylogeny revealed that it follows a similar pattern to *adhE* (Figure S1), being also statistically dependent of the niche (phylogenetically informed model testing, *p-value* = 1.0398e^-07^ Chi-squared test).

The other top predictive features are involved in multiple functions including carbohydrate metabolism (K01443, K08483, K01838), amino acid metabolism (K03402), osmotic stress (K03499, K03498, K04063), and DNA repair (K03111, K01669) (Figure 2B). Among these, we found *ptsI* (K08483), which was previously described to be absent in some FLAB species (16).

To test whether loss of *adhE* reduced the accuracy of RF classification, we removed the *adhE* feature from the dataset and reran the RF classifier. We obtained the same accuracy and similar precision (six false negatives and seven false positives with *adhE* vs seven false negatives vs six false positives without *adhE*; Table S3).

We also hypothesized that since bee-associated species largely stem from four clades, phylogenetic signal could be driving the RF classification. Several pieces of evidence suggest that phylogenetic relatedness is not the only driver of this classification. Firstly, we obtained accurate classifications across the entire phylogenetic spectrum both for multiple distantly related entire clades, but also for single species (i.e., *Holzapfelia floricola)* (Figure 1). This suggests that, apart from the common features shared by species belonging to monophyletic clades, some of the features are also shared with distantly related single species that are clustered within non-bee-associated clades. Secondly, looking at the most important KEGG annotations for the RF classifier (Figure 2B, Figure 2C), we could observe that many are absent from more than 75% of the species, suggesting that the same losses happened in multiple distantly related clades, independently. This indicates that irrespective of the phylogenetic relatedness, the same featured independently evolved in, at least, more than one clade.

We also looked at the species that were incorrectly classified (Table S3). Some of these may be explained by inaccuracies in our ecological categorization, which was based on the substrate of isolation of the strain for which the genome sequence was obtained. For instance, the *Apilactobacillus kunkeei* strain used in this work, which was identified as a false negative by the RF classifier (Figure S1, Table S3), was isolated from grape wine, but most strains of this species are known to be associated and isolated from the bee environment (14, 16, 20, 22).

### Metabolism related genes were largely lost in the MRCA of bee-associated LAB clades

To gain insight into functions that were most likely associated with the ecological shift, we next asked which genes were likely lost in the branches leading to bee-associated clades.

Hence, we looked for functions that were likely lost specifically in the MRCA of bee-associated clades and were therefore lost concomitantly with the ecological shift. Losses were inferred under a Wagner parsimony method [22], which assumes an unknown ancestral state and allows both transitions from 0 (loss) to 1 (gain) and the reverse (32). KEGG functional annotations for all species and the species tree topology (Figure 1) were used. This analysis revealed that significant losses took place in the MRCA of many of these clades, supporting previous evidence of genome reduction. For instance, under this model, 61 losses were inferred in the MRCA of bee-associated *Fructilactobacillus* - *Apilactobacillus* genera (Figure 3). Most losses across the four clades involved genes related with the metabolism of carbohydrates, amino acids, lipids and co-factors and vitamins (Figure 3). Importantly, we found that from the top 20 most import features for the RF classifier (Figure 2B), 13 were inferred to have been lost in the MRCA of at least one bee-associated clade (Table S3). Five out of the 13 genes were inferred to have been lost in the MRCA of two clades. Specifically, the two most important genes for the RF classifier, *adhE* and *ohyA*, were lost in the MRCA of two bee-associated clades, independently. For instance, loss of *AdhE* was inferred in the MRCA of both *Lactobacillus* and *Fructobacillus* (Table S3). Loss of *ptsI* gene (K08483) was also inferred in the MRCA of *Fructobacillus* and in the MRCA of *Fructilactobacillus* - *Apilactobacillus.* Two additional genes were inferred to have been independently lost in the branches leading to two distantly related bee-associated clades (K01443 and K01838). The remaining eight out of the 13 genes were inferred to have been lost in the MRCA of one clade only. These genes are however absent in many species belonging to other distantly related clades, suggesting that losses can occur after the ecological shift, in a stepwise manner. The remaining seven genes that are among the top 20 most important features for the random forest classifier were not found to have been lost in the MRCA of bee associated clades, suggesting that they were probably lost at distinct time points (either before or after the ecological shift) or present a patchy distribution.

**Figure 3.**
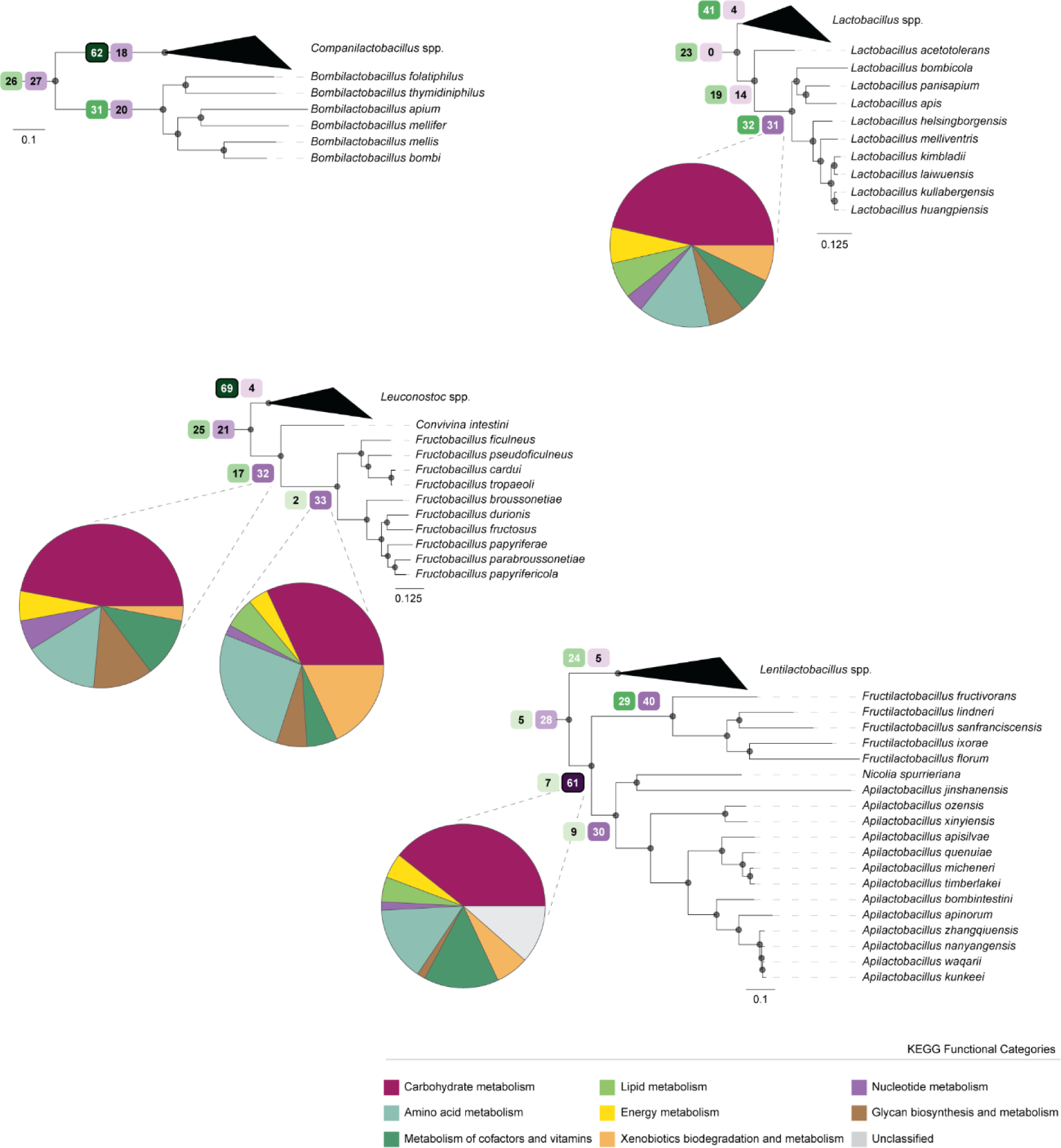
**Major losses of genes related to metabolic functions of occurred in the MRCA of bee associated clades**. Pruned trees of the four main groups of bee-associated species and closest relatives showing the number of annotated KEGG functional categories that were inferred to have been gained (green tones) or lost (purple tones) in the specific nodes corresponding to the MRCA of bee-associated species. Pie charts reflect the proportion of lost KEGG functional categories (annotated in KEGG Mapper) as indicated in the key. For *Bombilactobacillus* <5 KEGG were assigned a functional category.

We next inspected whether there was a statistically significant difference in the presence/absence pattern of certain biological functions between bee-associated species and their closest relatives. To ascertain this, we performed COG (Cluster of Orthologous Groups) analyses by comparing bee-associated species with their closest relatives (Figure 4). For all cases, bee-associated clades had lower numbers of proteins annotated in all the categories inspected, except for *Lactobacillus* spp. for which no differences were found (Figure S3, Table S4). For 13 out of the 26 COG categories, at least two bee-associated clades showed significant differences from their closest relatives. Some of the categories that showed significant differences were K (transcription), G (carbohydrate transport and metabolism), I (lipid transport and metabolism), and P (inorganic ion transport and metabolism). Loss of carbohydrate transport and metabolism functions was previously described for species belonging to the FLAB group (16, 21, 33), and it is in line with our aforementioned results. These losses seem to be more evident in *Fructobacillus* and *Fructilactobacillus* - *Apilactobacillus* clades (Figure 4). In the *Bombilactobacillus* clade, there was also a decrease in the number of genes assigned to category G (carbohydrate transport and metabolism) when compared to sister genus *Companilactobacillus*, however this difference was not statistically significant. Lipid transport and metabolism (I) is also statistically significantly less represented in bee-associated clades (Figure 4).

**Figure 4.**
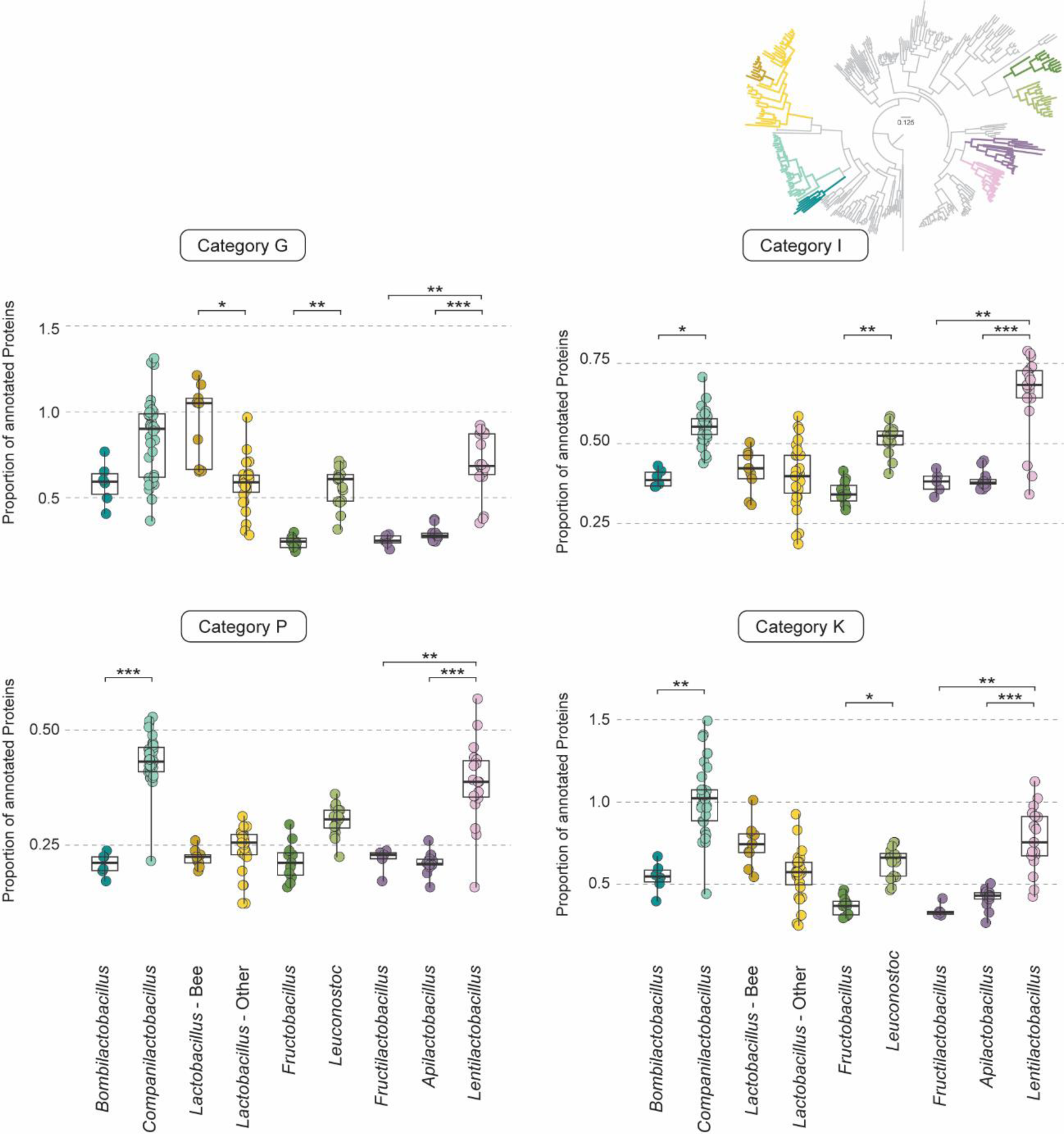
**Most bee associated clades have significantly fewer genes involved in carbohydrate and lipid transport and metabolism**. Boxplots show the proportion of annotated proteins in categories G (carbohydrate transport and metabolism), I (lipid transport and metabolism), P (inorganic ion transport and metabolism) and K (transcription) for bee-associated and their respective sister clades.

## Discussion

### Reductive evolution might be a hallmark of bee-associated microbes

One of the distinctive phenotypic traits found in FLAB is the preference for fructose over glucose (fructophily) (20). This unusual metabolic trait is hypothesized to have evolved in a background of lack of alcoholic fermentation and subsequent rewiring of sugar metabolism towards an alternative type of fermentation that involves fructose and mannitol as substrate and product, respectively (18, 20, 22, 25). We found that one of the most important genomic features for the RF classifier was the absence of Adh (alcohol dehydrogenase) activity (total or partial absence of the encoding *adhE* gene), which is the main determinant for fructophily in FLAB (19). Phenotypic characterization regarding sugar preference is unfortunately only available for a handful of species under study, however for species in which fructophily has been observed, absence of Adh activity is a hallmark (Figure S1) (13, 19, 21). While many other genes were lost in bee-associated bacterial species when compared to their counterparts that occupy distinct ecological niches, indicating that reductive evolution is pervasive in this ecological setting, specifically the loss of alcoholic fermentation-related genes seems to be strongly associated with other microorganisms thriving in the floral environment. This is supported by the multiple independent instances of loss of *adhE* across bee-associated LAB, but also by the loss of the entire alcoholic fermentation pathway in a fructophilic floral yeast clade (*Wickerhamiella/Starmerella* clade, Kingdom Fungi) (34, 35). Many of the species composing the *Wickerhamiella*/*Starmerella* yeast clade are intimately associated with bees and are many times isolated together with FLAB, from the same samples (36). Interestingly, the loss of alcoholic fermentation in this yeast clade is also seemingly associated with the emergence of fructophily, possibly also linked to a major metabolic rewiring towards the alternative mannitol fermentation pathway that uses fructose (and not glucose) as a substrate (37).

Similarly to FLAB, this yeast lineage also lost many metabolic abilities (35, 38, 39) supporting the notion that reduction of metabolic abilities might be pervasive across microorganisms adapting to the bee environment. Whether the set of genes and functions lost were similar among bee-associated yeasts and bacteria awaits further investigations.

### Were the losses adaptive or neutral byproducts of the switch to a new environment?

Gene loss is an important genetic mechanism contributing to phenotypic variation across the tree of life (38, 40–43). Gene losses can be adaptive in the sense that they are required for adaptation to a new environment, but they can also occur in a neutral fashion, as a consequence of relaxed selection following adaptation. If the timepoint in evolution when these losses occurred coincides with a period during which adaptation evolved, these two scenarios are difficult to distinguish without experimental validation, however we can draw some inferences based on the putative functions of the identified lost genes. Using complementary approaches involving machine learning and evolutionary analysis of homolog family sizes across LAB, we found that most predicted lost functions are related with metabolism. More specifically, we inferred that some of these losses might have occurred concomitantly with the adaptation period, meaning that they can be either adaptive or neutral. Although experimental validation of the phenotypic consequences of inactivation of these genes in a distinct genetic background would be required to distinguish between these two hypotheses, their functions (mainly sugar, lipid, and amino acid metabolism) suggest that losses were most likely the result of relaxed selection.

Interestingly, we find not only the same functions, but the same genes were sometimes lost independently across bee-associated LAB. While the recurrent evolution of the same trait under similar selective pressures across distantly related species can reflect the power of selection, it can also occur for reasons other than selection on that trait. If sources of variation are biased and only allow a limited number of changes, distantly related species may easily evolve convergent traits by non-adaptive processes owing to relaxed selection (44), constraints [5], mutational biases [11], or shared genetic variation [12]. Genetic constraints in particular (e.g. shared pleiotropic or epistatic effects) can be important drivers of patterns of convergence. For instance, if occupation of the bee-associated environment leads to the evolution of faster fructose consumption through loss of alcoholic fermentation, then other fermentation-related features may also evolve convergently as a consequence, although they are directly not under selection.

### The potential of statistical learning approaches for detecting fingerprints of adaptation

Machine learning algorithms generally make predictions about unknown data based on training and learning with known data. However, detecting the patterns in known data that make a certain feature of interest predictable can be extremely valuable to answer scientific questions, especially when those questions are better answered when analysing voluminous amounts and distinct types of data. Machine learning is an emerging powerful approach that has been used, for instance, to uncover the molecular bases of certain metabolic functions (29, 30, 45, 46) or to find phenotypic and genomic signatures of convergent evolution of certain ecological traits in yeasts (28). Here we reinforce the power of this approach by uncovering common gene loss events associated with adaptation to the bee environment. We not only found genes that have been previously associated with FLAB bacteria and linked to particular phenotypes (*adhE* – fructophily), but we uncovered new ones (e.g., *ohyA*) that have been recurrently and independently lost across bee-associated LAB. Overall, these results underscore how convergent evolutionary paths can be followed by microorganisms thriving in similar ecological contexts. Moreover, it underscores the power of machine learning to uncover fingerprints of convergent evolution, opening new and efficient avenues for the study of ecological adaptation.

## Materials and methods

### Genome annotation and phylogenetic inference

A total of 369 representative genomes of the family *Lactobacillaceae* and two outgroups (*Lactococcus lactis* and *Enterococcus massiliensis*) were retrieved from the NCBI (19 January 2023). For all species the complete proteome was predicted with AUGUSTUS v3.3.3 (47) using the complete gene model and *Staphylococcus aureus* as reference.

The species phylogeny was reconstructed with single copy orthogroups (SCO) present in at least 50% of the species obtained using OrthoFinder v2.5.4 (23) from the AUGUSTUS predicted proteomes. The SCO were aligned independently using MAFFT v7.407 (48) and then concatenated using a python script (https://github.com/santiagosnchez/ConcatFasta). The concatenated aligned file composed of 180 SCO was then used to infer a Maximum Likelihood (ML) phylogeny using IQ-TREE 1.6.11 (49) with a partition flag (-spp), an automatic detection of the best-fitting model and 1,000 ultrafast bootstrap replicates (50). The phylogeny was rooted using *Lactococcus lactis* and *Enterococcus massiliensis* (21).

### HMM-based search of *adhE* and *ohyA* across LAB

To search for *adhE* and *ohyA* across all strains, first a HMM profile was constructed. For that the reviewed protein of AdhE for *Escherichia coli* was recovered from UniProt (P0A9Q7) and OhyA for *Lactobacillus amylolyticus* (WP_127345688) were used as query for a BLASTp search in NCBI refseq database. Hits with an e-value lower than 0.001 were retrieved, up to a maximum of 100 hits. The sequences recovered were then aligned with MAFFT v7.407 (48) and a HMM profile for each gene was constructed in HMMER v3.3.3 (51). The HMM profile was used to score presence/absence of each gene and the respective copy number using Orthofisher v1.0.5 (26) with default parameters. Detected partial sequences of AdhE were subsequently analysed through BLASTp on NCBI to assessed which domain of the bifunctional protein was missing (aldehyde dehydrogenase or alcohol dehydrogenase).

### Inference of gene losses

Functional genomic annotations were performed using KofamScan [6] against KEGG database (ver. 2022-12-31) with default settings. To minimize false-positive assignments, taxon-specific profile databases were used, in this case prokaryotic proteins were annotated using ‘prokaryote.hal’. Proteins, for which the KofamScan did not yield a functional annotation (no K number assigned) were not reported in the output (’--no-report-unannotated’). Species for which a low number of proteins were mapped to a functional KEGG annotation were excluded from posterior analysis. To define the exclusion threshold a ratio between the number of coding sequences and annotated KEGG was determined for each species. Species with a ratio above 3 were excluded (Figure S2).

To infer losses across our dataset we used two approaches. The first, implied using the tool COGclassifier (52). COGclassifier assigns prokaryote protein sequences into COG (Cluster of Orthologous Genes) functional categories. This analysis was performed for all strains in bee-associated clades and their respective sister clades. The proportion of proteins assigned to each category was calculated by dividing the number of annotated proteins from each strain to the total number of annotated proteins for a given category.

To infer gene losses in specific nodes we used a parsimony-based approach implemented in Count (53), under the Wagner parsimony method. The number of losses were inferred in the MRCA of the bee-associated clades and respective sister clades and the KO annotations involved were retrieved and subsequently analysed with KEGGmapper to determine their functional categories. In both cases KEGG annotations generated by KofamScan were used as input.

### Machine Learning

To test whether we could predict the floral-bee niche from KEGG Orthology genomic data, we used a random forest algorithm. We trained a machine learning algorithm built by an XGBoost (1.7.3) random forest classifier (XGBRFClassifier()) with the parameters “ max_depth=12 and n_estimators=100; all other parameters were in their default settings. The max_depth parameter specifies the depth of each decision tree, determining how complex the random forest will be to prevent overfitting while maintaining accuracy. The n_estimators parameter specifies the number of decision trees in the forest—after testing the increase in accuracy while increasing each of these parameters, we found that having a higher max_depth or more decision trees per random forest did not further increase accuracy.

The random forest algorithm was trained on 90% of the data, and used the remaining 10% for cross-validation, using the RepeatedStratifiedKFold and cross_val_score functions from the sklearn.model_selection (1.2.1) package. Cross validation is a method for assessing accuracy involving 10 trials, each of which holds back a random 10% of the training data for testing (54, 55) Given the unbalanced nature of the dataset, we used balanced accuracy, which takes the mean of the true positive rate and the true negative rate, since there were unequal numbers of growers and non-growers in many of these substrates. For both measures, an accuracy value of 50% would be equivalent to randomly guessing. Top features were automatically generated by the XGBRFClassifier function using Gini importance, which uses node impurity (the amount of variance in environmental niche for strains that either have or do not have this KEGG Ortholog).

### Statistical analysis

All statistical analyses were performed in RStudio. All datasets were firstly tested for normality with Shapiro-Wilk normality test (shapiro.test). Graphs and statistics were done with the package ggstatsplot (56) where for each case it was specified the type of analysis (parametric or non-parametric) according to Shapiro-Wilk results. For all statistical tests, ‘***’ corresponds to a *p-*value of 0.001, ‘**’ *p-*value 0.01, and ‘*’ *p-*value 0.05.

For phylogeny-informed statistical analyses, namely differences in genomes size, protein-coding genes and GC content between different niches (bee-associated and other) a phylogenetic ANOVA using the function ‘phylANOVA’ implemented in the R package phytools (24, 57) was used.

To assess if there was a dependence relationship between niche (bee / other) and AdhE (presence / absence) when accounting for phylogeny, we conducted model testing of four hypothesis using phytools v 2.1.1 (57). Specifically, we tested if (i) the niche pattern and the occurrence of AdhE is independent of one another (null hypothesis), (ii) the niche pattern is dependent of AdhE, (iii) the pattern of occurrence of AdhE is dependent on niche, and (iv) the patterns of niche and AdhE are interdependent. We evaluated model fit using weighted Akaike information criterion and compared the best fitting model to the null hypothesis using a Chi-squared test. We also did the same test for the OhyA protein which has a similar distribution as AdhE. The hypotheses tested were the same.

## Acknowledgements and Funding

We thank members of the Yeast Genomic Lab for helpful discussions, specifically Paula Gonçalves for constructive suggestions to the manuscript. We also thank Jacob L. Steenwyk for helpful suggestions on statistical methodologies.

This work was financed by national funds from FCT—Fundação para a Ciência e a Tecnologia, I.P. (FCT/MCTES; https://www.fct.pt/) in the scope of project UIDB/04378/2020 and UIDP/04378/2020 of the Research Unit on Applied Molecular Biosciences— UCIBIO and project LA/P/0140/2020 of the Associate Laboratory Institute for Health and Bioeconomy— i4HB. It was further supported by grants PTDC/BIA-EVL/0604/2021 (to CG), from FCT/MCTES. Computational work was carried out with support of INCD funded by FCT and FEDER under the project 01/SAICT/2016 n° 022153 and the grants 2023.09581.CPCA.A1 (to AP). Research in the AR lab is supported by the National Science Foundation (DEB-2110404), the NIH/National Institute of Allergy and Infectious Diseases (R01 AI153356), and the Burroughs Wellcome Fund.

## Competing interests

AR is a scientific consultant for LifeMine Therapeutics, Inc.

## Figures and Tables

**Figure S1.**
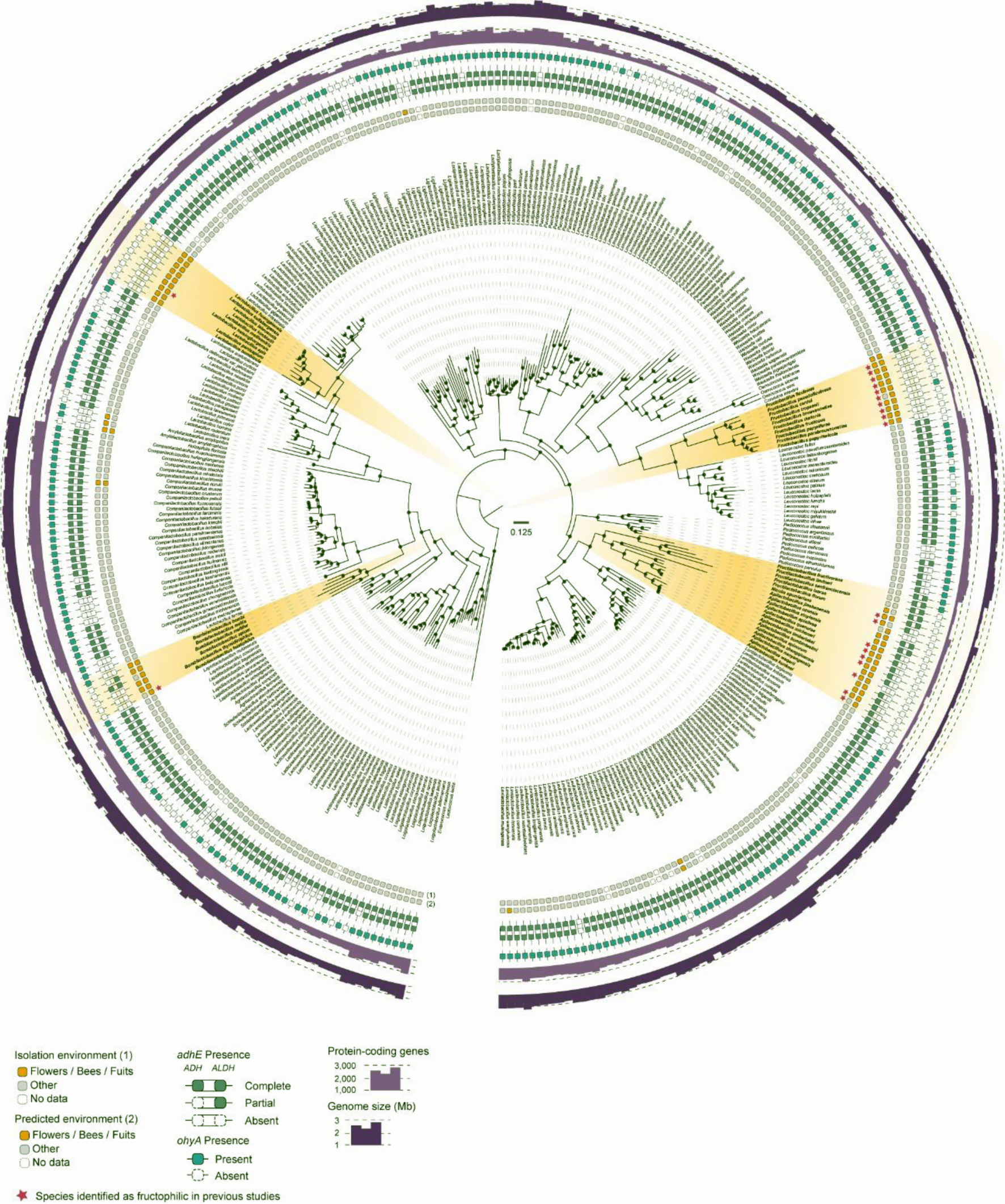
Maximum likelihood phylogenomic tree comprising 369 Lactobacillaceae species inferred from the concatenated alignment of 180 SCO and rooted with *Lactococcus lactis* and *Enterococcus massiliensis*. The isolation source and predicted environment are depicted in the first two rings, respectively. Presence/absence of *adhE* and *ohyA*, number of protein-coding genes and genome size are also shown.

**Figure S2.**
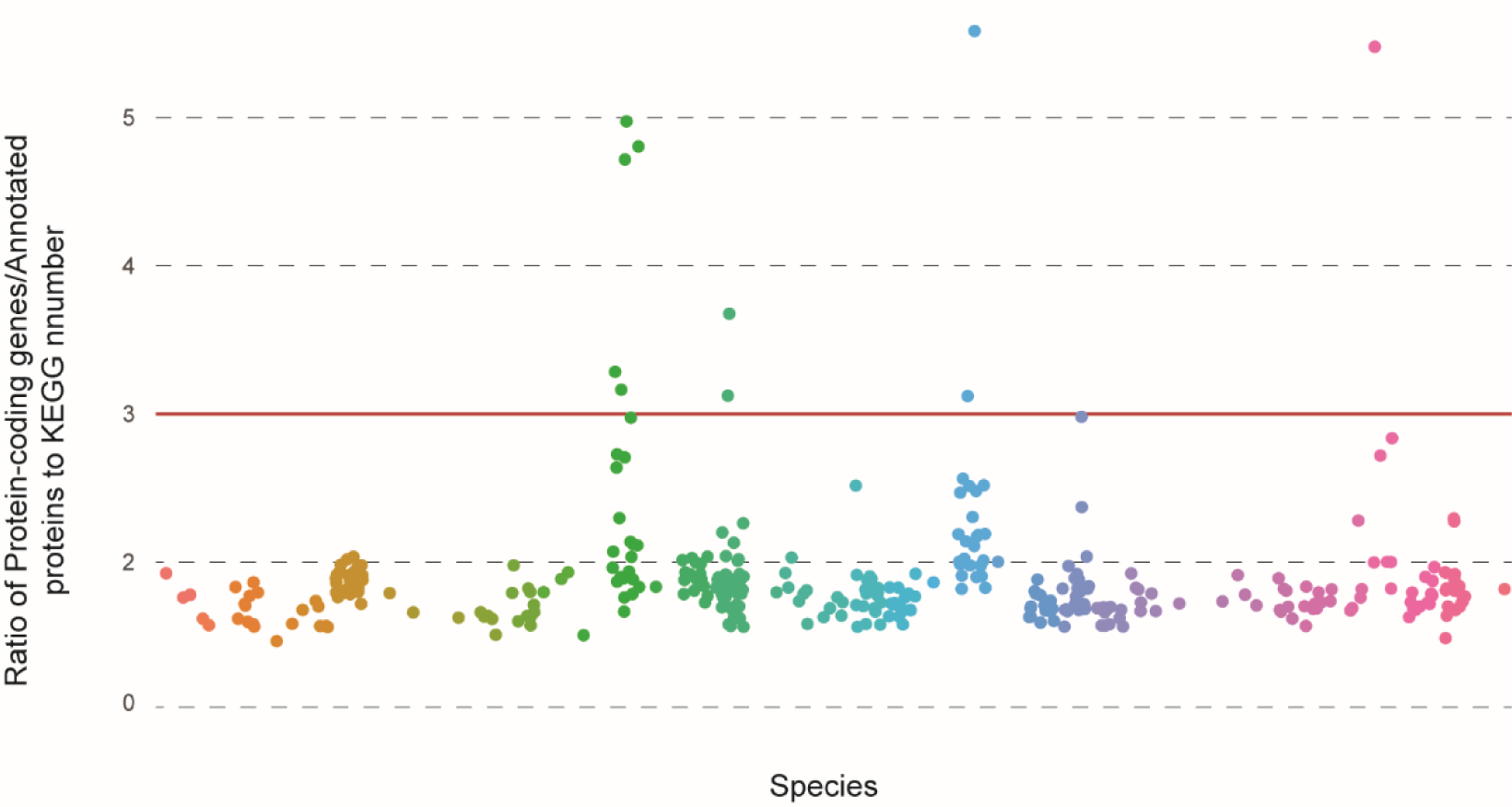
Ratio between the number of protein-coding genes and the number of annotated proteins to a KEGG number (KO) for each species. The different genera are given by different colours. The threshold applied (ratio = 3) is marked by a red line.

**Figure S3.**
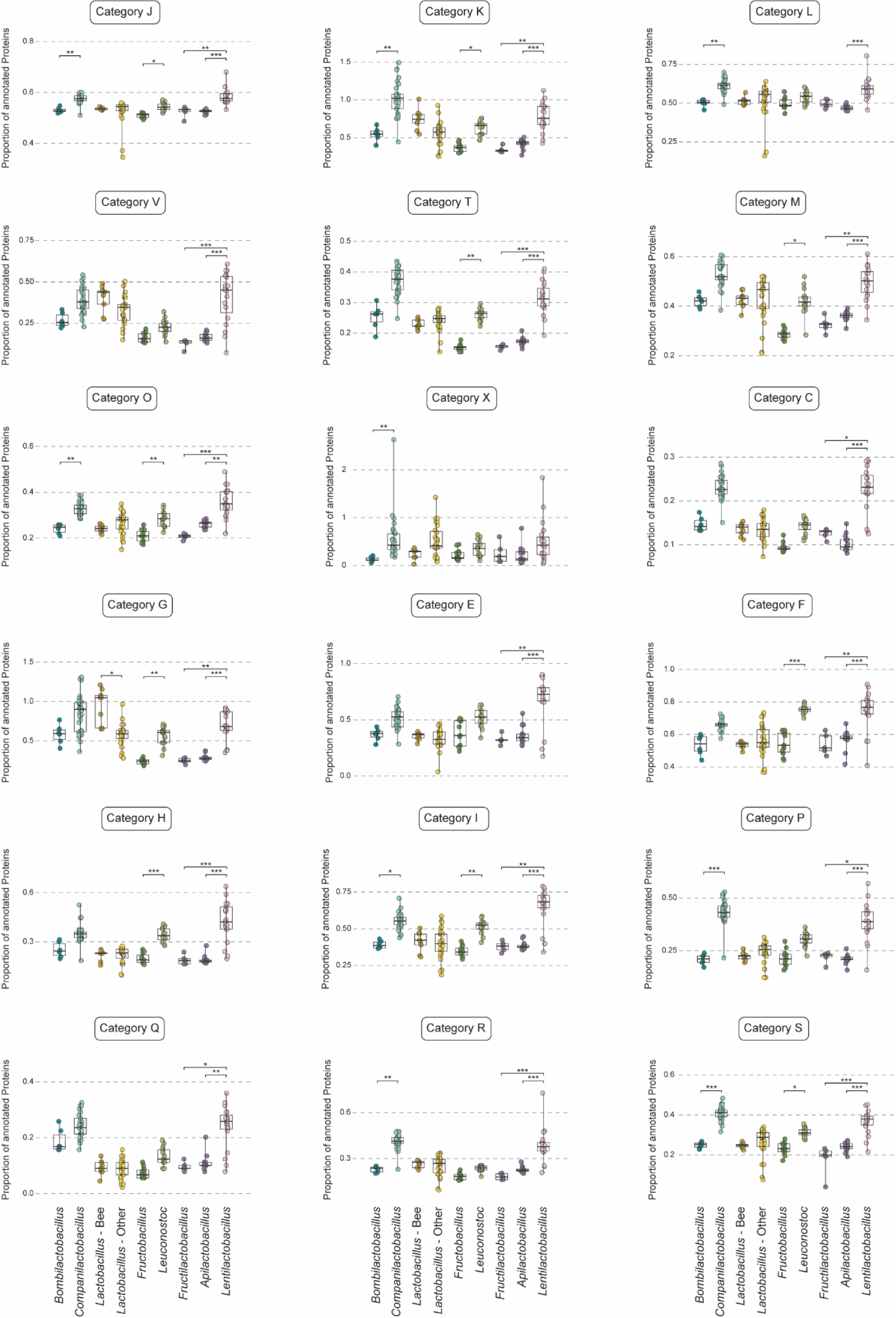
Boxplots show the proportion of annotated proteins in COG categories for bee-associated and their respective sister clades.

Table S1 - List of Lactobacillaceae species used in this work for the reconstruction of the phylogenomic tree and subsequent genomic analysis.

Table S2 – Tables containing information about the machine learning results, namely incorrectly identified species, top 20 important features and the proportion of bee-associated species and others across the top 20 features.

Table S3 – List of KEGG numbers gained, lost, contracted, and expanded in the four main clades of bee-associated species, given by Counts.

Table S4 – List of the annotated proteins by COG category for the clades of bee-associated species and respective sister clades. Data used for Figure 4 and Figure S2.

## References

1. Moran NA. 2002. Microbial Minimalism: Genome Reduction in Bacterial Pathogens. Cell 108:583–586.

2. Keeling PJ, Fast NM. 2002. Microsporidia: biology and evolution of highly reduced intracellular parasites. Annu Rev Microbiol 56:93–116.

3. Wadi L, Reinke AW. 2020. Evolution of microsporidia: An extremely successful group of eukaryotic intracellular parasites. PLOS Pathogens 16:e1008276.

4. Corradi N. 2015. Microsporidia: Eukaryotic Intracellular Parasites Shaped by Gene Loss and Horizontal Gene Transfers. Annu Rev Microbiol 69:167–83.

5. Pombert JF, Haag KL, Beidas S, Ebert D, Keeling PJ. 2015. The Ordospora colligata genome: Evolution of extreme reduction in microsporidia and host-to-parasite horizontal gene transfer. mBio 6.

6. McCutcheon JP, Moran NA. 2012. Extreme genome reduction in symbiotic bacteria. Nature Reviews Microbiology 10:13–26.

7. Murray GGR, Charlesworth J, Miller EL, Casey MJ, Lloyd CT, Gottschalk M, Tucker AW, Welch JJ, Weinert LA. 2021. Genome Reduction Is Associated with Bacterial Pathogenicity across Different Scales of Temporal and Ecological Divergence. Molecular Biology and Evolution 38:1570–1579.

8. Bobay L-M, Ochman H. 2018. Factors driving effective population size and pan-genome evolution in bacteria. BMC Evolutionary Biology 18:153.

9. Vasquez YM, Bennett GM. 2022. A complex interplay of evolutionary forces continues to shape ancient co-occurring symbiont genomes. iScience 25:104786.

10. Moran NA. 1996. Accelerated evolution and Muller’s rachet in endosymbiotic bacteria. Proc Natl Acad Sci U S A 93:2873–8.

11. Hottes AK, Freddolino PL, Khare A, Donnell ZN, Liu JC, Tavazoie S. 2013. Bacterial adaptation through loss of function. PLoS Genet 9:e1003617.

12. Sokurenko EV, Hasty DL, Dykhuizen DE. 1999. Pathoadaptive mutations: gene loss and variation in bacterial pathogens. Trends Microbiol 7:191–5.

13. Filannino P, Di Cagno R, Tlais AZA, Cantatore V, Gobbetti M. 2019. Fructose-rich niches traced the evolution of lactic acid bacteria toward fructophilic species. Crit Rev Microbiol 45:65–81.

14. Endo A, Salminen S. 2013. Honeybees and beehives are rich sources for fructophilic lactic acid bacteria. Syst Appl Microbiol 36:444–8.

15. Endo A, Futagawa-Endo Y, Dicks LM. 2009. Isolation and characterization of fructophilic lactic acid bacteria from fructose-rich niches. Syst Appl Microbiol 32:593–600.

16. Maeno S, Tanizawa Y, Kanesaki Y, Kubota E, Kumar H, Dicks L, Salminen S, Nakagawa J, Arita M, Endo A. 2016. Genomic characterization of a fructophilic bee symbiont Lactobacillus kunkeei reveals its niche-specific adaptation. Syst Appl Microbiol 39:516–526.

17. Salman SM, GhadahmohammedSaleh. Fructophilic lactic acid bacteria symbionts in honeybees – a key role to antimicrobial activities, p. *In* (ed),

18. Filannino P, Di Cagno R, Addante R, Pontonio E, Gobbetti M. 2016. Metabolism of Fructophilic Lactic Acid Bacteria Isolated from the genus-species Apis mellifera L. Bee Gut: Phenolic Acids as External Electron Acceptors. Applied and Environmental Microbiology 82:6899.

19. Maeno S, Kajikawa A, Dicks L, Endo A. 2019. Introduction of bifunctional alcohol/acetaldehyde dehydrogenase gene (adhE) in Fructobacillus fructosus settled its fructophilic characteristics. Res Microbiol 170:35–42.

20. Endo A, Maeno S, Tanizawa Y, Kneifel W, Arita M, Dicks L, Salminen S. 2018. Fructophilic Lactic Acid Bacteria, a Unique Group of Fructose-Fermenting Microbes. Appl Environ Microbiol 84.

21. Maeno S, Nishimura H, Tanizawa Y, Dicks L, Arita M, Endo A. 2021. Unique niche- specific adaptation of fructophilic lactic acid bacteria and proposal of three Apilactobacillus species as novel members of the group. BMC Microbiology 21:41.

22. Endo A, Tanizawa Y, Tanaka N, Maeno S, Kumar H, Shiwa Y, Okada S, Yoshikawa H, Dicks L, Nakagawa J, Arita M. 2015. Comparative genomics of Fructobacillus spp. and Leuconostoc spp. reveals niche-specific evolution of Fructobacillus spp. BMC Genomics 16:1117.

23. Emms DM, Kelly S. 2019. OrthoFinder: phylogenetic orthology inference for comparative genomics. Genome Biology 20:238.

24. Adams DC, Collyer ML. 2018. Phylogenetic ANOVA: Group-clade aggregation, biological challenges, and a refined permutation procedure. Evolution 72:1204–1215.

25. Endo A, Tanaka N, Oikawa Y, Okada S, Dicks L. 2014. Fructophilic Characteristics of Fructobacillus spp. may be due to the Absence of an Alcohol/Acetaldehyde Dehydrogenase Gene (adhE). Current Microbiology 68:531–535.

26. Steenwyk JL, Rokas A. 2021. orthofisher: a broadly applicable tool for automated gene identification and retrieval. G3 Genes|Genomes|Genetics 11:jkab250.

27. Behare PV, Mazhar S, Pennone V, McAuliffe O. 2020. Evaluation of lactic acid bacteria strains isolated from fructose-rich environments for their mannitol-production and milk-gelation abilities. Journal of Dairy Science 103:11138–11151.

28. Gonçalves C, Harrison MC, Steenwyk JL, Opulente DA, LaBella AL, Wolters JF, Zhou X, Shen XX, Groenewald M, Hittinger CT, Rokas A. 2023. Diverse signatures of convergent evolution in cacti-associated yeasts. bioRxiv doi:10.1101/2023.09.14.557833.

29. Opulente DA, LaBella AL, Harrison MC, Wolters JF, Liu C, Li Y, Kominek J, Steenwyk JL, Stoneman HR, VanDenAvond J, Miller CR, Langdon QK, Silva M, Gonçalves C, Ubbelohde EJ, Li Y, Buh KV, Jarzyna M, Haase MAB, Rosa CA, Čadež N, Libkind D, DeVirgilio JH, Hulfachor AB, Kurtzman CP, Sampaio JP, Gonçalves P, Zhou X, Shen XX, Groenewald M, Rokas A, Hittinger CT. 2024. Genomic factors shape carbon and nitrogen metabolic niche breadth across Saccharomycotina yeasts. Science 384.

30. Harrison M-C, Ubbelohde EJ, LaBella AL, Opulente DA, Wolters JF, Zhou X, Shen X-X, Groenewald M, Hittinger CT, Rokas A. 2023. Machine learning enables identification of an alternative yeast galactose utilization pathway. PNAS 121.

31. Radka CD, Batte JL, Frank MW, Young BM, Rock CO. 2021. Structure and mechanism of Staphylococcus aureus oleate hydratase (OhyA). J Biol Chem 296:100252.

32. Alekseyenko AV, Lee CJ, Suchard MA. 2008. Wagner and Dollo: a stochastic duet by composing two parsimonious solos. Syst Biol 57:772–84.

33. Endo A, Maeno S, Tanizawa Y, Kneifel W, Arita M, Dicks L, Salminen S. 2018. Fructophilic Lactic Acid Bacteria, a Unique Group of Fructose-Fermenting Microbes. Applied and environmental microbiology 84:e01290–18.

34. Gonçalves P, Gonçalves C, Brito PH, Sampaio JP. 2020. The Wickerhamiella/Starmerella clade—A treasure trove for the study of the evolution of yeast metabolism. Yeast n/a.

35. Gonçalves C, Wisecaver JH, Kominek J, Oom MS, Leandro MJ, Shen X-X, Opulente DA, Zhou X, Peris D, Kurtzman CP, Hittinger CT, Rokas A, Gonçalves P. 2018. Evidence for loss and reacquisition of alcoholic fermentation in a fructophilic yeast lineage. eLife 7:e33034.

36. de Oliveira Scoaris D, Hughes FM, Silveira MA, Evans JD, Pettis JS, Bastos E, Rosa CA. 2021. Microbial communities associated with honey bees in Brazil and in the United States. Braz J Microbiol 52:2097–2115.

37. Gonçalves C, Ferreira C, Gonçalves LG, Turner DL, Leandro MJ, Salema-Oom M, Santos H, Gonçalves P. 2019. A New Pathway for Mannitol Metabolism in Yeasts Suggests a Link to the Evolution of Alcoholic Fermentation. Front Microbiol 10:2510.

38. Shen XX, Opulente DA, Kominek J, Zhou X, Steenwyk JL, Buh KV, Haase MAB, Wisecaver JH, Wang M, Doering DT, Boudouris JT, Schneider RM, Langdon QK, Ohkuma M, Endoh R, Takashima M, Manabe RI, Cadez N, Libkind D, Rosa CA, DeVirgilio J, Hulfachor AB, Groenewald M, Kurtzman CP, Hittinger CT, Rokas A. 2018. Tempo and Mode of Genome Evolution in the Budding Yeast Subphylum. Cell 175:1533–1545.e20.

39. Gonçalves C, Gonçalves P. 2019. Multilayered horizontal operon transfers from bacteria reconstruct a thiamine salvage pathway in yeasts. Proc Natl Acad Sci U S A 116:22219–22228.

40. Olson MV. 1999. When less is more: gene loss as an engine of evolutionary change. American journal of human genetics 64:18–23.

41. Albalat R, Cañestro C. 2016. Evolution by gene loss. Nature Reviews Genetics 17:379–391.

42. Guijarro-Clarke C, Holland PWH, Paps J. 2020. Widespread patterns of gene loss in the evolution of the animal kingdom. Nature Ecology & Evolution 4:519–523.

43. Bhattacharya D, Qiu H, Lee J, Su Yoon H, Weber APM, Price DC. 2018. When Less is More: Red Algae as Models for Studying Gene Loss and Genome Evolution in Eukaryotes. Critical Reviews in Plant Sciences 37:81–99.

44. McCutcheon JP, McDonald BR, Moran NA. 2009. Convergent evolution of metabolic roles in bacterial co-symbionts of insects. Proceedings of the National Academy of Sciences 106:15394–15399.

45. Nalabothu RL, Fisher KJ, LaBella AL, Meyer TA, Opulente DA, Wolters JF, Rokas A, Hittinger CT. 2023. Codon Optimization Improves the Prediction of Xylose Metabolism from Gene Content in Budding Yeasts. Molecular Biology and Evolution 40.

46. Olivia R, Allison SW, Antonis R. 2023. Predicting fungal secondary metabolite activity from biosynthetic gene cluster data using machine learning. bioRxiv doi:10.1101/2023.09.12.557468:2023.09.12.557468.

47. Stanke M, Keller O, Gunduz I, Hayes A, Waack S, Morgenstern B. 2006. AUGUSTUS: ab initio prediction of alternative transcripts. Nucleic acids research 34:W435–W439.

48. Katoh K, Standley DM. 2014. MAFFT: iterative refinement and additional methods. Methods Mol Biol 1079:131–46.

49. Nguyen LT, Schmidt HA, von Haeseler A, Minh BQ. 2015. IQ-TREE: a fast and effective stochastic algorithm for estimating maximum-likelihood phylogenies. Mol Biol Evol 32:268–74.

50. Hoang DT, Chernomor O, von Haeseler A, Minh BQ, Vinh LS. 2018. UFBoot2: Improving the Ultrafast Bootstrap Approximation. Mol Biol Evol 35:518–522.

51. Finn RD, Clements J, Eddy SR. 2011. HMMER web server: interactive sequence similarity searching. Nucleic Acids Res 39:W29–37.

52. Tatusov RL, Galperin MY, Natale DA, Koonin EV. 2000. The COG database: a tool for genome-scale analysis of protein functions and evolution. Nucleic Acids Res 28:33–6.

53. Csűös M. 2010. Count: evolutionary analysis of phylogenetic profiles with parsimony and likelihood. Bioinformatics 26:1910–1912.

54. Pedregosa F, Varoquaux G, Gramfort A, Michel V, Thirion B, Grisel O, Blondel M, Prettenhofer P, Weiss R, Dubourg V, Vanderplas J, Passos A, Cournapeau D, Brucher M, Perrot M, Duchesnay É. 2011. Scikit-learn: Machine Learning in Python. J Mach Learn Res 12:2825–2830.

55. Chen T, Guestrin C. 2016. XGBoost: A Scalable Tree Boosting System, abstr Proceedings of the 22nd ACM SIGKDD International Conference on Knowledge Discovery and Data Mining, San Francisco, California, USA, Association for Computing Machinery,

56. Patil I. 2021. Visualizations with statistical details: The ‘ggstatsplot’ approach. The Journal of Open Source Software 6:3167.

57. Revell LJ. 2012. phytools: an R package for phylogenetic comparative biology (and other things). Methods in Ecology and Evolution 3:217–223.

